# Rapid genetic diversification of *Bacteroides thetaiotaomicron* in mono-associated mice revealed through deep population-level sequencing

**DOI:** 10.1101/2025.06.24.661302

**Authors:** Christos Zioutis, Michaela Lang, Fatima Pereira, Olga Bochkareva, Ekaterina Kolodyazhnaya, Jay Osvatic, Kathy D. McCoy, Sven Künzel, Hann Fokt, John F. Baines, David Berry

## Abstract

Bacteria often feature short generation times and large populations, thereby allowing them to quickly evolve and adapt to new environments. Although it is known that gut bacteria can evolve on relatively short time scales, the extent of genetic diversification of bacteria in the gut environment remains underexplored. Here, we characterize the genetic diversification of the gut commensal *Bacteroides thetaiotaomicron* during 28 days of colonization of germ-free mice using deep shotgun sequencing as well as genome analysis of evolved isolates. We detect thousands of genetic polymorphisms as early as three days post inoculation and observe highly dynamic genetic diversity in the distal gut. We identify multiple haplotypes of a phase-variable polysaccharide utilization locus (*BT2260* - *BT2268*) and propose that phase variation may be an important mechanism for diversification and adaptation in the gut. In addition, we find evidence that hybrid two-component system (*HTCS)* regulators are mutational hotspots. We identify multiple persistent and parallelly evolved genetic polymorphisms in genes, including the TonB-dependent transporter *BT0867* - a homolog of *BF3581* from the commensal colonization factor (*ccf*) in *B. fragilis*. Lastly, we find that the small intestine accumulated approximately 20 times more polymorphisms compared to the large intestine, highlighting overall the importance of studying spatiotemporal distribution of genetic variants. These results underscore the prevalence of rapid genetic diversification of gut bacteria, which may have important implications for adaptation as well as interactions in the microbiome and with the host.

**Importance:** Studying the within-host evolution of gut commensals is an essential step for understanding the role of microbiome in health and disease. It can provide insights into the mechanisms underlying the development of various gastrointestinal disorders, metabolic conditions, autoimmune diseases, and other health disorders. Additionally, this kind of research can further drive the development of personalized therapies, such as strain-level or gene specific interventions for improving health outcomes. Here, we report extensive genetic variation within days upon colonization of mice with *B. thetaiotaomicron* and identify genes that accumulate persistent and highly prevalent genetic polymorphisms across a mouse population. We also detect several haplotypes in phase-variable loci. Altogether, our findings underscore the rapid pace of genetic diversification and phase variation upon colonization of the gut environment.

## Introduction

The gastrointestinal tract is densely colonized by microorganisms that play important roles in host nutrition and physiology (1–3). Ecological disturbances in the gut microbiome can have profound effects on health (4–6). In addition to changes over macroevolutionary timescales (7), there is evidence that continued microbial evolution occurs even within the lifetime of individual hosts (8–11), highlighting the importance of adaptability to environmental disturbances on ecologically relevant timescales. Bacteria have large population sizes as well as short generation times, which are ideal characteristics for rapid genetic diversification (12, 13). Random point mutations such as single nucleotide polymorphisms are widely considered to be important primers for phenotypic change (14, 15). However, structural variants such as large inversions can also be drivers for genetic and phenotypic heterogeneity.

Phase variation describes genetic and epigenetic mechanisms that are involved in heritable, but reversible phenotypic variability, and result in local increases in genetic variation (16). Remarkably, stochastic fluctuation between phenotypes over a small number of generations has been mathematically modeled to be the most beneficial strategy in fluctuating environments (17, 18). In *Bacteroides*, phase variation has been implicated in immune system evasion and resistance to bacteriophage infection (19–25).

*Bacteroides* are an abundant and key genus in the human gut microbiota (26). They specialize in the degradation of polysaccharides derived both from the host (27) and the diet (28–30), which is reflected in the presence of various polysaccharide utilization loci (PULs) in their genomes (31, 32). These discrete genetic structures contain all necessary genes for glycan degradation, including glycoside hydrolases with cleaving specificities for different polysaccharides (33). Pivotal to the pathway are the TonB-dependent *SusC* family proteins that form a trans-membrane complex with *SusD* family proteins (34, 35), through which oligosaccharides are imported to the periplasm for further degradation. Additionally, transcriptional regulators sense periplasmic oligosaccharide concentration and can trigger a positive feedback loop (36). In recent years, prominent species from this genus have been established as model organisms for genetic engineering (37) and colonization experiments (11), which have led to the identification of colonization factors, such as the *ccf* locus in *B. fragilis* (38) and other *in vivo* fitness determinants in *B. thetaiotaomicron* (39, 40). Nevertheless, much research has been focused on tracking the abundances of genetically introduced transposon mutants. In this work, we employ an experimental evolution approach using mice monocolonized with *B. thetaiotaomicron* VPI-5482 (41) to investigate the characteristics and dynamics of genetic diversification in the gut ecosystem.

## Results

### Phase-variable loci are active during colonization

We monitored the appearance of genetic polymorphisms in *B. thetaiotaomicron VPI- 5482* at multiple time points during the first 28 days of gut colonization of germ-free mice in two separate evolution experiments. Our approach was based on deep shotgun sequencing of feces and intestinal contents as well as genome sequencing of fecal isolates (Fig. 1). The initial analysis of genetic variants in the sequenced populations revealed patterns of highly correlated frequency trajectories for some polymorphisms, which we named haplotype-like signatures (HLSs). Although HLSs could be attributed to genetic hitchhiking after the emergence of multiple dominant haplotypes, it could also be the result of multiple genomic inversions in phase-variable regions. As it would be impracticable to discriminate those scenarios strictly from the population data, we evaluated phase variation in genomes from 180 isolates retrieved from fecal samples (∼30 per d28 fecal sample from Facility A). Phase-variable regions were defined as genetic loci where at least one recombination event that introduced gene shuffling was observed in the genome.

**FIG 1.**
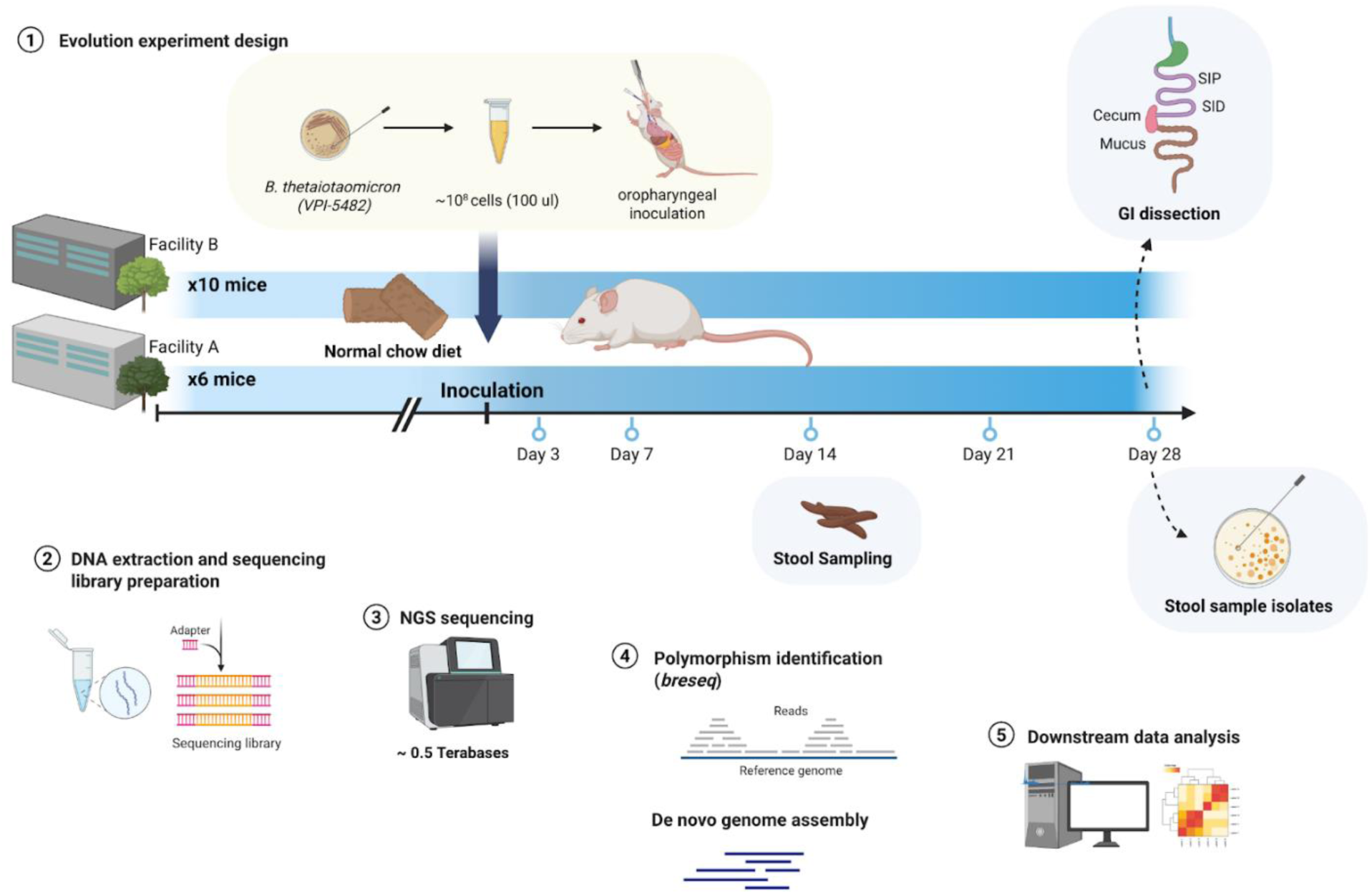
Overview of the experimental design and sequencing analysis workflow. (**1**) Germ-free mice were orally inoculated with B. thetaiotaomicron and fecal pellets were collected over a period of 28 days in two separate experiments (Facilities A and B). On the last day of the experiment, mice were dissected and contents from other intestinal compartments were collected (Facility B; proximal small intestine [SIP], distal small intestine [SID], cecum, mucus). Single colonies were also isolated from fecal samples (Facility A). (**2**) DNA was extracted and multi-indexed sequencing libraries were prepared for (**3**) Whole genome sequencing (WGS), yielding ∼0.5 terabases of sequences. (**4**) For population samples, polymorphisms were identified in comparison with the reference genome with breseq, while genomes from single colonies were de-novo assembled, before any (**5**) downstream analysis.

We detected seven active phase-variable regions (also referred to as shufflons) in genomes of gut-evolved *B. thetaiotaomicron* (Fig. 2). Three of these shufflons were involved in modification of *susC* family genes, two affected restriction endonucleases, and two only altered gene orientation. Apart from the already described *BT1040- BT1046* locus (24), we were able to experimentally validate phase haplotypes for *BT2260*, *BT2264* and *BT2268* genes in predicted PUL 27 and PUL 30, which has been previously speculated to be a shufflon due to the similarly positioned integrase (34). Despite the similar PUL-like structure in these two loci, they encode different integrases (*BT1041* and *BT2267* have ∼20% aa identity) as well as *susCD* gene pairs. Genomic repeats found on the borders of inversions (49 nucleotides in *BT1040*-*BT1046* and 22 nucleotides of in *BT2260*-*BT2268)* do not share any motifs, which support the hypothesis that phase variation in these loci is moderated by different recombinases. In turn, the *BT2267* is homologous to tyrosine site-specific recombinase (Tsr0667, ∼80% identity), which was shown to moderate numerous shufflon-type DNA inversions in loci that contain or are colocalized with *susCD*-family genes in *Bacteroides fragilis* (23). The inverted repeat *AGTTCTGCAAAGACTTTGCAGA* found in the *BT2260*-*BT2268* locus is in agreement with the motif sequences observed in Tsr0667-regulated invertible regions (23). Additionally, we observed phase variation coupled with modifications of two *SusC* paralogs *BT3239*-*BT3240* (aa identity ∼80%) in PUL 52, which lacks the centrally located integrase (Fig. 2), but has also been predicted to contain inverted repeats (IRs) involved in phase variation (21). A common feature of phase variation in these loci is the relocation of the N-terminal region of the two *susC* family genes, which contains a signal peptide (Sec/SPI) followed by a carboxypeptidase D regulatory like domain (pfam: *PF13715*), and in some cases is interleaved by a signal transduction domain (pfam: *PF7660*) (34).

**FIG 2.**
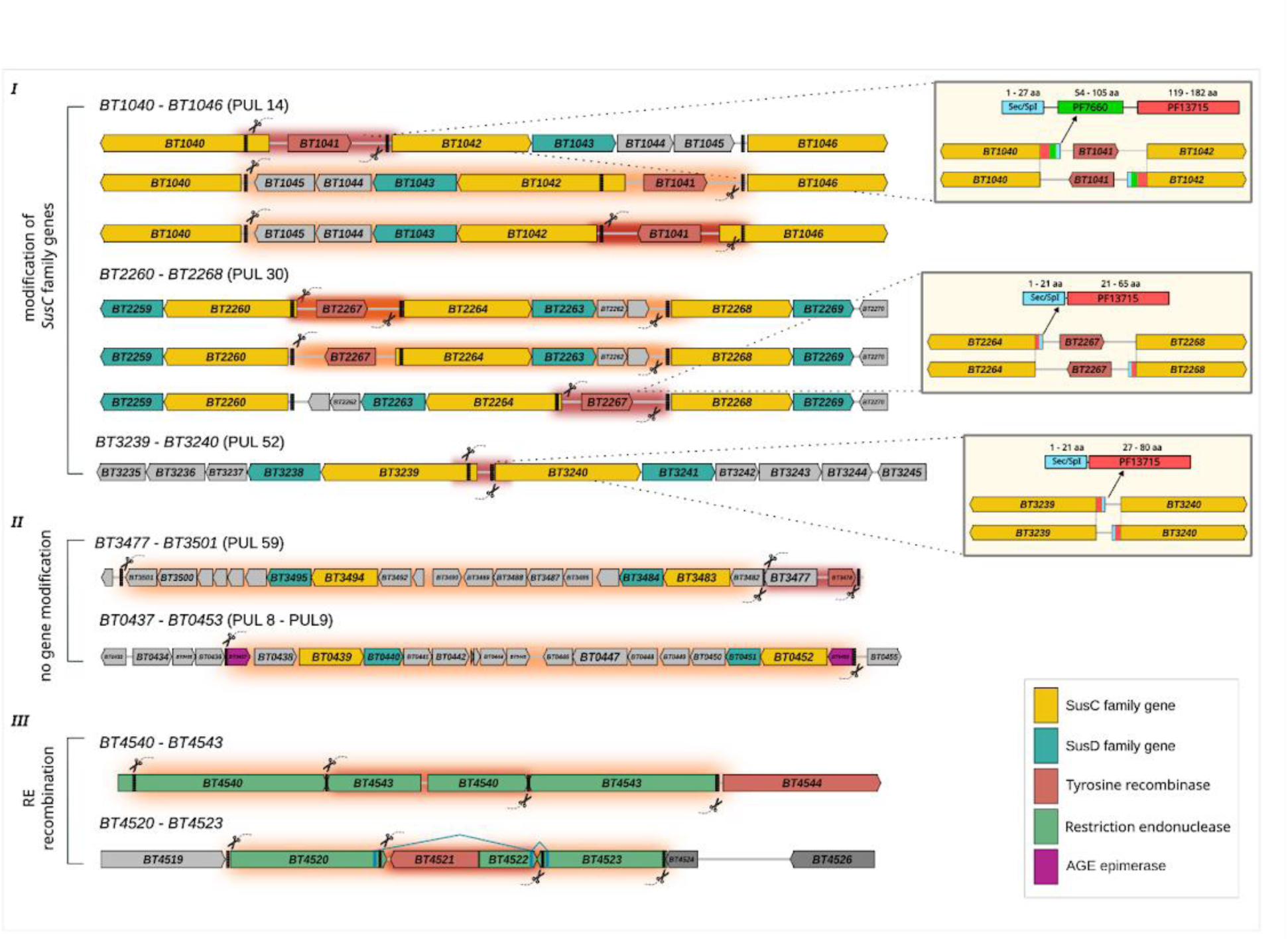
Phase haplotypes detected in multiple loci. Three types of phase variation were observed; I) within PULs with susC family genes modification, II) within PULs but without modification of susC family genes and III) associated with restriction endonucleases (RE) recombination. Zoom in plots for susC family genes modifications, indicate pfam protein domains and predicted signal peptides, which are relocated in the N-terminal region across haplotypes.

The phase haplotypes in *BT0437*-*BT0453* (PUL 8 - PUL 9) and *BT3477*-*BT3501* (PUL 59) shufflons did not result in any changes in the gene sequences apart from their respective orientation (Fig. 2), although we did not investigate whether these rearrangements could play a role in promoter sequences or transcriptional activity. Lastly, the *BT4540*-*BT4543* and *BT4520*-*BT4523* shufflons encoded four and three homologs of restriction endonucleases, respectively, which were inverted and recombined to create multiple variants (Fig. 2).

Analysis of 19 publicly-available complete *B. thetaiotaomicron* genomes revealed phase haplotypes in five loci (Fig. S1). *BT1040*-*BT1046* and *BT2260*-*BT2268* PUL-related phase-variable loci were conserved across all complete assemblies, possibly indicating their importance for *B. thetaiotaomicron* fitness, while others were disrupted by large insertions or deletions. The distribution of phase haplotypes did not display phylogenetic signal across *B. thetaiotaomicron* strains in any of the loci, highlighting the frequency of these events. These results emphasize the potential role of phase variation as a mechanism for *B. thetaiotaomicron’s* rapid genetic diversification in the gut.

### *B. thetaiotaomicron* undergoes rapid and extensive genetic diversification

In addition to phase-variable loci, we detected overall 63,657 unique *de novo* genetic polymorphisms in deeply-sequenced (median genome coverage ∼ 200X) fecal samples. The vast majority of detected variants were single nucleotide polymorphisms (SNPs), with deletions (DEL) and insertions (INS) constituting a minor fraction (∼8-10%; Fig. 3A). High polymorphism levels were apparent as early as three days post-inoculation. We observed a much higher number of polymorphisms at d3 in some mice (median = 2800, CI = [1000,11000]; Wilcoxon paired test of: d3-d14, p = 0.008, d3-d21, p = 0.013). However, overall levels did not change considerably after one week of colonization, fluctuating around a median of ∼1K polymorphisms (Wilcoxon paired test, n.s.; Fig. 3A). A very low level of standing genetic variation was present in the expanded clonal ancestral populations, but did not contribute substantially to this early spike at d3 (Facility A ∼ 98.2%, Facility B ∼ 99.9% of polymorphisms were detected only *in vivo*). Although many of the SNPs were located in intergenic regions, non-synonymous mutations accumulated at higher numbers compared to synonymous mutations (ratio 95% CI [3.3 - 3.7], Wilcoxon paired test, d3-d28, p < 0.001) throughout the experiments (Fig. 3B). Mean nucleotide diversity (π) in intragenic regions was similar after the first week (Wilcoxon test, n.s.; Fig. 3C), while polymorphisms affected about 1% of the genome. Despite statistical similarity at the cohort level, individual trajectories were characterized by oscillations in genetic diversity. This result indicates that mutations can rapidly emerge in considerable numbers and abundances (>= 1% relative frequency), but also be quickly lost from the population.

**FIG 3.**
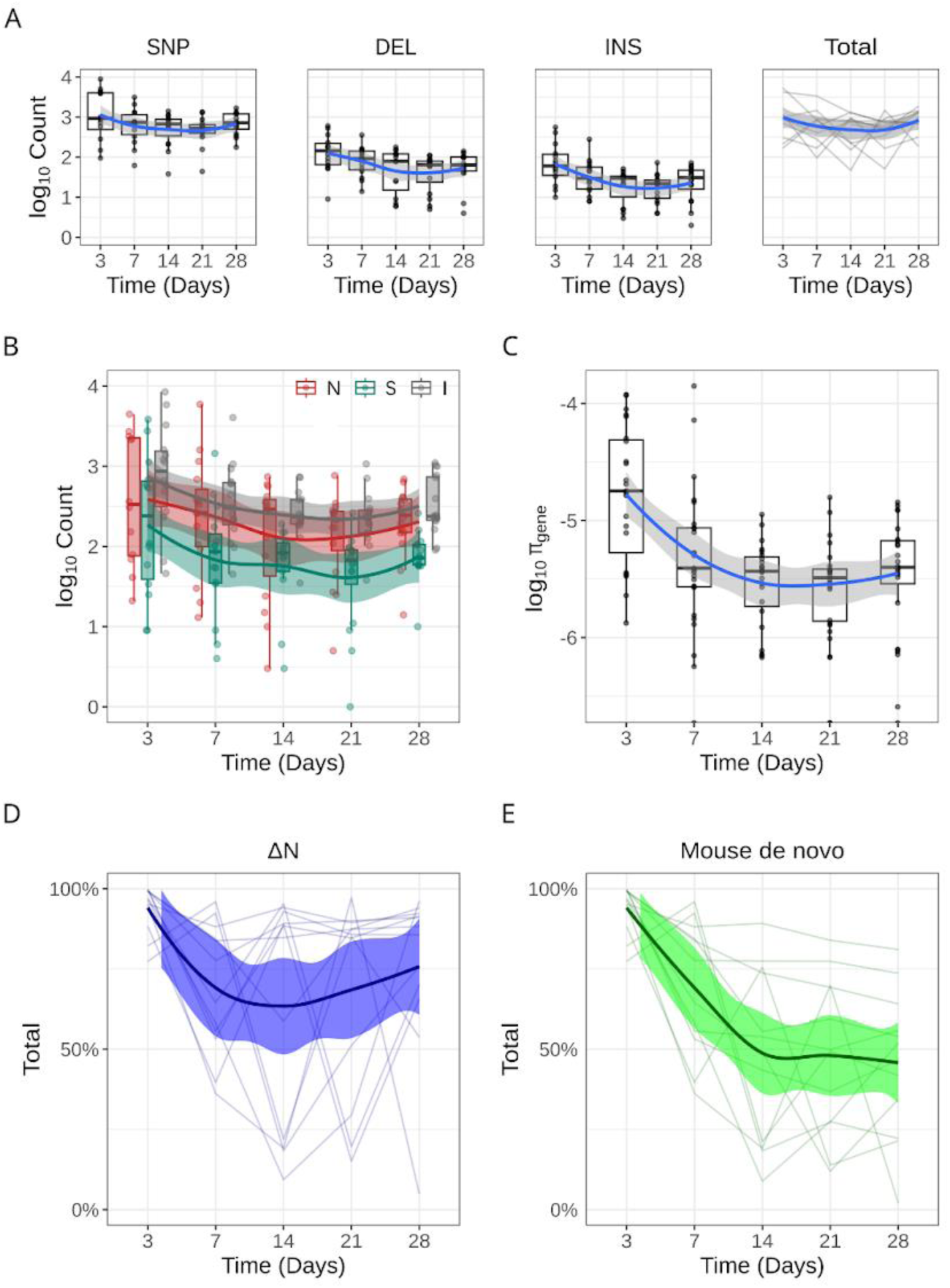
High levels of standing genetic variation are detected early and persist throughout the experiment, despite volatile dynamics. **(A)** Log10-transformed counts of single nucleotide polymorphisms (SNP), deletions (DEL), insertions (INS) and total number of polymorphisms over 28 days of in vivo evolution. **(B)** Log10-transformed counts of non-synonymous polymorphisms (N) in red, synonymous (S) in blue and intergenic (I) in grey over time in days. **(C)** Log10-transformed intragenic nucleotide diversity (π) over time in days. **(D)** Genetic variation sampled in each time point if not detected in the exact previous time point as a percentage of the total variation in the time point (ΔN) over time for each mouse. **(E)** De novo genetic variation emerged in each mouse evolutionary trajectory as a percentage of the total variation in each timepoint over time.

Although mice are prone to microbial transmission through feces due to their coprophagic behavior, co-housing did not significantly influence the observed genetic variability according to non-parametric multivariate analysis of variance (PERMANOVA, p > 0.05). Similarly, there was no statistical support for mouse-driven variability. Approximately 9% of the genetic variability was attributable to the time point (PERMANOVA; Time point: r^2^ = 0.09, p = 0.001), while ∼6% of the genetic variability was associated with the individual experiments. (PERMANOVA; Facility: r^2^ = 0.059, p = 0.001). We next evaluated polymorphism dynamics by quantifying two measures at each time point; the proportion of polymorphisms that were not detected in the previous time point (ΔN), and the proportion of polymorphisms that were detected for the first time in a mouse trajectory (mouse *de novo*). We found that despite the total number of polymorphisms in the population, there was a continuous detection of *de novo* mutations that could replace up to 97% of the variation within 7 days (Fig. 3D). Interestingly, the mean ΔN did not decline after the first week of colonization (Wilcoxon test, d7-d28, ns), in contrast to the mouse *de novo* fraction, which decreased over time (Pearson, r =-0.32, p = 0.01; Fig. 3E). These outcomes, combined with fluctuations in the total number of polymorphisms, underscore the highly dynamic genetic landscape in this ecosystem.

### Hotspots and parallelism in genetic diversification

In order to evaluate whether certain genes or functions were hotspots for genetic diversification, we first examined the distribution of polymorphisms across different functional groups at three hierarchical levels: COG categories, COG IDs and genes. Although there was no statistical enrichment in polymorphisms for specific COG categories and IDs, we found multiple genes that accumulated a significantly higher number of polymorphisms, not only in individual samples, but generally across the dataset (Fig. 4A). The majority of identified genes encoded for hybrid two-component system (HTCS) proteins, which frequently act as regulators of PUL expression. Indeed, we find that these highly mutated regulator genes are associated with PULs involved in both host-derived/mucin glycan utilization, such as O-glycans (PUL 32, PUL 38, PUL 76) (27) and heparin/heparan sulfate (PUL 85) (30, 42), as well as diet-derived polysaccharides such as rhamnogalacturonan I (PUL 77), rhamnogalacturonan II (PUL 92), homogalacturonan (PUL 75), arabinan (PUL 7), arabinogalactan (PUL 65) (29), and α-mannan (PUL 90) (43).

**FIG 4.**
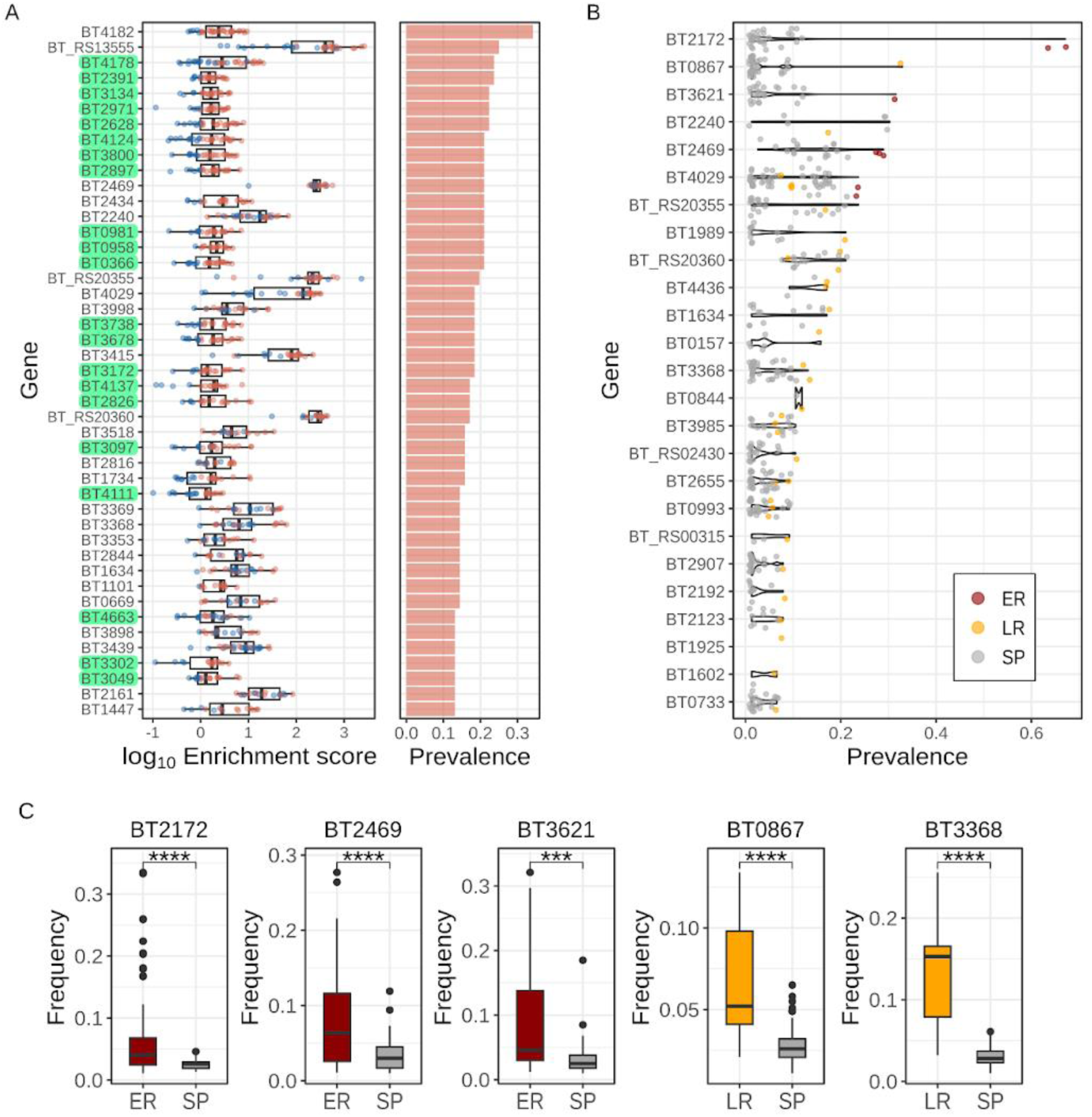
Hotspots of mutations and parallel genetic diversification in evolved populations. **(A)** Log10-transformed enrichment score for all genes that appear in our dataset at least 10 times with enrichment score > 1 and at least 5 polymorphisms. Genes with enrichment score > 1 and at least 5 polymorphisms in a sample are shown in red (left). Dataset prevalence of cases with enrichment score > 1 and 5 polymorphisms for each gene (right). *HTCS* genes are highlighted in green. **(B)** On the y-axis, genes with at least one early and/or late response polymorphism. Early response polymorphisms (ER), in orange, persist for at least four time points in three evolutionary trajectories at minimum. Late response polymorphisms (LR), in red, are detected at the last two timepoints (d21 and d28 or d28) across three evolutionary trajectories at minimum. The rest, in grey, are considered sporadic polymorphisms (SP). On the x-axis, dataset prevalence for each polymorphism. **(C)** Comparison of population frequency between ER or LR polymorphisms and SP for *BT2172*, *BT2469*, *BT0967*, *BT3368* genes respectively from left to right.

We next evaluated genes with polymorphisms that either appeared early across multiple mice and persisted throughout the experiment (Early response - ER) or that appeared later in the experiment across multiple mice (Late response - LR) (Fig. 4B). Polymorphisms in a TonB-dependent transporter (TBDT) gene located in PUL27 (*BT2172*) showed the highest persistence across multiple mice. Other genes with ER mutations included an integrase (*BT2469*), an amidase enhancer (*BT3621*), and multiple hypothetical proteins. Another TBDT gene, *BT0867*, acquired a LR mutation (T765S) within a porin superfamily domain (SSF56935). This gene is located in PUL 12 and has 86% identity on protein level to the *BF3581* of the commensal colonization factor genetic locus (*ccf*) previously characterized in *B. fragilis* (38, 44). LR mutations were also identified in genes not associated with PUL structures, including a glycosyltransferase (*BT3368*), a TPR domain containing protein (*BT2240*), a beta-galactosidase (*BT0993*), and other hypothetical proteins. We identified cases of genes where these putatively adaptive mutations were not only detected in considerable frequency in the population, but also at a much higher level compared to sporadic mutations on the same gene (Fig. 4C), which is suggestive of a fitness advantage.

### Distinct patterns of genetic diversification in small and large intestines

Collection of samples from different intestinal locations - the colonic mucus, cecum, proximal small intestine (SIP), and distal small intestine (SID) - helped us to evaluate the influence of intestinal microenvironments on genetic diversification. We found that genetic diversity between the small (SI) and large (LI) intestinal compartments had pronounced differences (Fig. 5A). These differences were heavily driven by the difference in total number of polymorphisms between these two groups (Wilcoxon test, p < 0.001; Fig. 5B). Remarkably, more than 120,000 different polymorphisms were unique to SI samples, and this number was more than 20 times higher than those detected exclusively in the large intestine as well as those common between the two compartments (Fig. 5C). To assess the association of polymorphisms to intestinal compartments, we used polymorphism abundances in a mixed effects model. No genetic variants unique to the LI were significantly associated, due to their lower overall prevalence (<0.2). On the other hand, mutations with the highest coefficients in the model were unique to the SI (Fig. 5C). Many of these were located within genes related to the cell wall/membrane structure (Fig. 5D), such as a cell surface protein (*BT1771*), a putative glycosyltransferase TagX (*BT3622*), a LPS biosynthesis protein (*BT0392*) and a fibrillin family protein (*BT1014*). Other SI specific targets of diversification were involved in nutrient uptake (heme chaperone *BT2168*, secreted endoglycosidase *BT1038*, permease *BT0834*), ribosome maturation (*BT3836*, *BT0913*) and DNA repair and modification (helicase *BT4603*, methyltransferase *BT4626*, gyrase *BT0899*). Collectively, our findings suggest that *B. thetaiotaomicron* rapidly diversifies to respond to the intestinal environment and effectively occupies available ecological niches.

**FIG 5.**
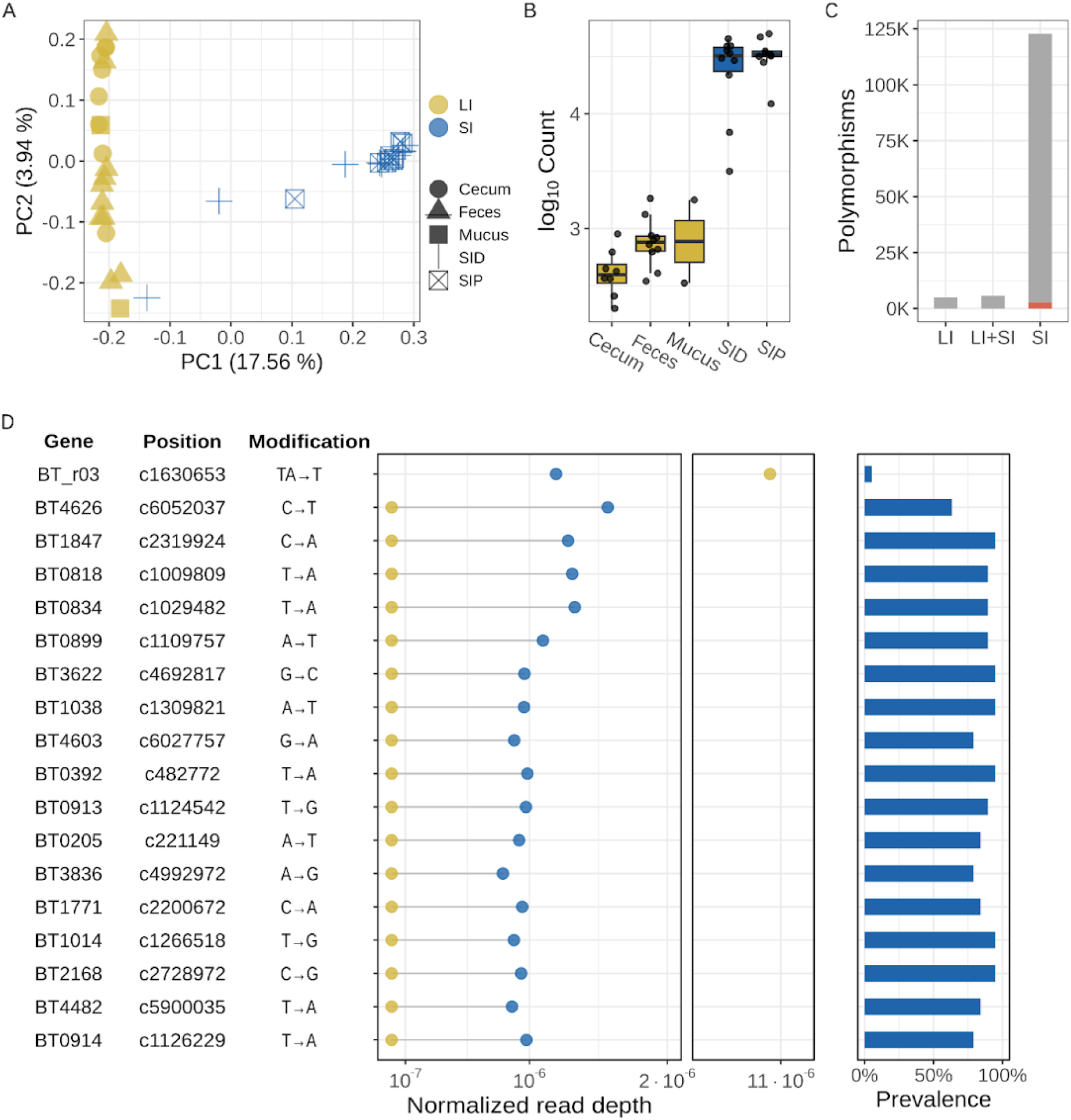
Patterns of genetic diversification in small and large intestine compartments. **(A)** Principal Coordinate Analysis (PCoA) on Bray-Curtis distances calculated from binary polymorphism abundance data. Samples from large intestine (LI) compartments are shown in yellow, whereas samples from small intestine (SI) are shown in blue. Different shapes indicate individual compartments; circle for Cecum, triangle for Feces, square for Mucus, cross for Small Intestine Proximal (SIP) and ballot box with X for Small Intestine Distal (SID). **(B)** Log10-transformed count of polymorphisms across all sampled intestinal compartments; Cecum, Feces, Mucus, SID, SIP from left to right. **(C)** Count of polymorphisms detected in samples from both large and small intestine (common), only in large intestine samples (LI) and only in small intestine samples (SI). In red, the fraction of significantly associated polymorphisms. **(D)** (left) On the y-axis, the 99th percentile of polymorphisms with the highest coefficient in absolute numbers, in our Multivariate Generalized Linear Model (GLM). On the x-axis, mean normalized read depth for small, in yellow and large, in blue, intestine samples. (right) Polymorphism prevalence in large intestine samples as percentage.

## Discussion

Experimental evolution is a powerful method to study microbial adaptation to a range of conditions. In this study, we report rapid dynamics in genetic diversification of *B. thetaiotaomicron* and potential implications for its adaptation to the intestinal environment. Phase variation in *Bacteroides* has primarily been described in CPS loci, and has been viewed as an important mechanism for modulating antigenic variability and phage evasion and thereby providing a colonization advantage (24, 25, 44). This phase variation is presumed to be mediated by the reversible inversion of promoters facilitated by site-specific integrases (19, 20). There have been other proposed phase-variable loci in *B. thetaiotaomicron* and *B. fragilis* based on genome scans for inverted repeats (21) including surface components and restriction modification proteins (22). Interestingly, the two restriction-modification loci found in the present study have also been reported in a recent study (45). Yet, to our knowledge, there is only one characterized locus in *B. thetaiotaomicron* where phase-variable inversions introduce changes in both gene orientation and the coding sequence, namely the *BT1040*-*BT1046* (PUL 14) shufflon (24). Here, we provide experimental evidence for phase variation in six additional genetic loci in *B. thetaiotaomicron*. We found that these recombination events can be mediated in several steps, yielding multiple phase haplotypes. *BT2268* and *BT2269* genes have been proposed as mucus-associated functional factors (MAFF) due to their IgA-mediated upregulation in mucus-resident bacteria (46), while transposon knock-out mutants of *BT2262*-*BT2264* showed a growth defect *in vivo* (47), suggesting an important role of this shufflon in gut colonization. The multiple phase haplotypes from the genomes of evolved *B. thetaiotaomicron* lineages indicate that these variants are present at a substantial frequency in the *in vivo* populations. It is noteworthy that these variants emerged as abundant in the population within 28 days of *in vivo* evolution, in line with the timescale in which phase-variable promoters in CPS loci change their expression levels (44) as well as the emergence of other phase-variable variants upon gut colonization (45). Collectively, these findings suggest that phase variation may be an important mechanism for adaptive diversification in the gut in addition to the classical CPS modulation strategy. The potential fitness advantage of such rearrangements as well as variation rates should be explored in future studies.

Cryptic variants and their contribution to adaptation are often overlooked, but our deep metagenomic sequencing approach allowed us to observe rapid and extensive genetic diversification of *B. thetaiotaomicron* upon gut colonization in germ-free mice. We detected thousands of *de novo* polymorphisms as soon as three days after inoculation (Fig. 3A). High levels of standing variation have been shown to be critical for the emergence of adaptive lineages in the gut of gnotobiotic mice while enhancing the effect of clonal interference (48). In fact, rapid adaptive diversification within a few days or weeks has been observed in colonization studies in *Bacteroides* spp. (40, 45, 49) and *Escherichia coli* (50–53). In the human microbiome, *de novo* genetic modifications are thought to serve as the main source of genetic diversity up to a 6-month timescale (9) and *Bacteroides* species retain high levels of standing variation (∼10^4^ SNPs), which may play an important role in fast adaptation to fitness landscape shifts driven by ecological disturbances (10, 11). Convergent evolution within host lineages and asymmetric distribution of adaptive mutations in different human populations further underscore the importance of *de novo* diversification (8). Clonal interference across multiple lineages with similar fitness effects has been demonstrated in colonization experiments (48, 50) and could explain the absence of high frequency polymorphisms in the populations.

We observed that genetic variation could be replaced entirely within seven days. This may be driven in part by the intensity of environmental disturbances. However, we speculate that polymorphisms shared across evolutionary trajectories can reflect underlying universal adaptive responses. Moreover, persistence within trajectories could indicate a strategy for a more long-term adaptation. The most persistent mutations were located in *BT2172* (LEU14-> [ARG14, ILE14]), a signal transduction TonB-dependent transporter (TBDT) in a PUL predicted to target host glycans that does not contain a *susD* paralog and thus potentially has a novel function (34). We also identified a late response mutation (T756S) in *BT0867*, the closest homolog of *BF3581* from the *ccf* locus discovered in *B. fragilis*. Mutations in this particular amino acid (T756I and T756A) have also been reported in a recent colonization study (45).

21 out of the 32 predicted hybrid two-component system (HTCS) genes (28) in *B. thetaiotaomicron* were hotspots for mutations with a high prevalence across samples. HTCS regulators constitute a key sensing mechanism in *Bacteroides* and are coupled with the regulation of PULs (28, 54–57). Deletion of *BT2391* has been previously shown to promote *B. thetaiotaomicron* colonization by increasing the ability of mucin glycan utilization and mucosal adhesion (58) while ΔBT3172 mutant strains have reduced fitness *in vivo* (54). In a recent functional genetic study, *BT4111* had increased fitness for growth in pectin, polygalacturonic acid and rhamnogalacturonan I, and a growth advantage for *BT4663* in heparin from porcine (47). Altogether, the higher accumulation of mutations in HTCS genes suggests an important evolutionary strategy for *B. thetaiotaomicron in vivo*.

Spatial stratification of bacteria cross-sectionally and along the intestinal tract has been well documented (38, 59). However, the evolutionary consequences of these spatial gradients are yet unexplored. Here, we observed significantly higher levels of genetic diversity in the small intestine compared to the large intestine. Higher growth rates, challenges from the physiological characteristics, and more frequent environmental disturbances (60–62) could enhance genetic diversification. We find that almost half of the polymorphisms observed in the large intestinal compartment were also detected in the small intestine. This could be attributed to the direction of the luminal fluid influx, as those polymorphisms could happen earlier in time and then translocate to more distal parts of the GI tract. While *Bacteroides* predominantly colonize the large intestine (63), members of the Bacteroidales have been detected in the murine ileum (64, 65). In the small intestine, bacteria digest simple carbohydrates and peptides (66) and produce vitamins (67). Higher oxygen levels, bile acid concentrations, and secreted antimicrobial peptides may constitute harsher environmental conditions, however the SI may also constitute an otherwise unavailable niche for *B. thetaiotomicron* in the mono-associated mouse model due to the absence of competing species. Possible adaptations to facilitate resilience and immune system invasion through changes in the cell wall/membrane as suggested by prevalent mutations in cell surface proteins (*BT1771*, *BT0392*), efficient nutrient uptake (*BT1038*; predicted secreted endoglycosidase, *BT0834*; permease) or even optimization of heme utilization (*BT2168*; predicted heme chaperone) (68), as well as sufficient responses to oxidative stress through DNA repair mechanisms (*BT4603*; helicase, *BT4626*; methyltransferase, *BT0899*; gyrase) could be vital for survival in this environment.

Future colonization experiments or *in vitro* fitness assays with mutant strains under controlled conditions are necessary to shed light into the underlying biological functions of mutations observed in the present study. *B. thetaiotaomicron* VPI-5482 is a human isolate, thus similar colonization experiments with mouse-specific strains could provide a better understanding of the adaptive landscape independent of the host. Additionally, studies with denser longitudinal sampling or lineage tracking can increase the resolution of the dynamics observed, provided that analysis of genetic diversity is not restricted to lineage frequencies but extends to include any *de novo* polymorphisms.

Our work presents evidence for multiple mechanisms of evolution of *B. thetaiotaomicron* and suggests that phase variation might be selected as a wide-spread adaptation strategy in this highly fluctuating environment, as it represents a larger fraction of the genetic variation landscape than previously assumed. Adaptability of microbial populations has numerous implications for community stability and resilience in the gut. Therefore, systematic study of intra-species diversity and their driving evolutionary processes can open new avenues to directed and personalized interventions in the gut microbiome.

## Materials and Methods

### Experimental design

We conducted two independent colonization experiments in germ-free mice at two separate facilities. The first experiment was conducted at the Clean Mouse Facility (CMF), University of Bern, Switzerland (Facility A) and the second was conducted at the Max Planck Institute for Evolutionary Biology, Plön, Germany (Facility B). Germ-free mice (Facility A, n=6; Facility B, n=10) were orally gavaged with 10^8^ cells of *B. thetaiotaomicron* VPI-5482. All mice were fed a normal chow diet. Mice at Facility A were fed autoclaved (132°C for 20 min) vitamin-fortified (3437, Kliba Nafag) chow. Mice at Facility B were fed sterilized 50 kGy V1124–927 Sniff (Soest, Deutschland) chow. Fecal pellets were collected at five time points (3, 7, 14, 21, and 28 days post inoculation). Additionally, 180 *B. thetaiotaomicron* colonies (30 per mouse) were isolated after growth of Facility A d28 fecal samples in BHI medium agar plates. At the end of the Facility B experiment, mice were dissected and luminal contents from the proximal small intestine (SIP), distal small intestine (SID), cecum, as well as the colonic mucus were collected additionally to the fecal pellets. For Facility A, all mouse experiments were performed in accordance with Swiss Federal regulation licensure. For Facility B, permission for animal husbandry is given by the Veterinäramt Kreis Plön (1401-144/PLÖ-004697) and for the experiment by the Ministerium für Landwirtschaft, ländliche Räume, Europa und Verbraucherschutz (MLLEV; permit 97-8/16).

### Bacterial strains and mouse inoculation

For mouse inoculation, *B. thetaiotaomicron* type strain VPI-5482 commercially available from DSMZ (DSM 2079), was grown in brain heart infusion (BHI) medium with supplements (BHIs, 37 g/l BHI, 5 g/l yeast extract, 1 g/l NaHCO_3_, 1 g/l L-cysteine, 1 mg/l vitamin K1, 5 mg/l hemin) in an anaerobic tent (Coy Labs, USA). This anaerobic culture was further diluted in Phosphate-Buffered Saline for a final concentration of 1 x 10^9^ cells/ ml and finally mice were orally gavaged with 100 µl (∼10^8^ cells).

### NGS library preparation and sequencing

For samples collected throughout the experiment, DNA was extracted from 25 mg of fecal pellet or 400 µl of intestinal content (for other sample types) with a standard CTAB, phenol/chloroform extraction method. For the isolates, we cultured glycerol-preserved fecal samples onto agar plates with BHIs medium under anaerobic conditions. Colonies were restreaked, to ensure clonality. Single colonies were grown overnight in 5 ml BHIs medium and from liquid cultures DNA was extracted using the DNeasy Blood and Tissue Kit (Qiagen), according to the manufacturer’s instructions. NGS libraries were prepared with *Nextera DNA XT* and *NebNext Ultra II FS* kits (Illumina). In the latter case, we followed manufacturer’s instructions for 350 bp insert size and DNA shearing was performed in a Covaris S220 Series instrument. DNA was quantified with Quant-IT Picogreen dsDNA and Qubit dsDNA HS assay kits, and libraries were multiplexed and pooled for each sequencing lane accordingly, so that an estimated genome coverage of ∼ 200X and ∼50X to be yielded for population samples and isolates respectively. Final fragment size distribution and concentration was carried on TapeStation 4100 and libraries were sequenced on the *Illumina HiSeq2500* (125 bp PE reads) platform at the Next Generation Sequencing facility at the Vienna BioCenter Core Facilities (VBCF) or *Illumina HiSeq3000* (150 bp PE reads) and *Illumina NovaSeq6000* (150 bp PE reads) at CeMM’s (Research Center for Molecular Medicine of the Austrian Academy of Sciences) biomedical sequencing facility (BSF). Overall, 122 population-wide samples were sequenced; two from the *B. thetaiotaomicron* inoculum culture, 80 from feces and 40 from intestinal compartments, as well as 180 isolates, that yielded in total 3.37 billion PE reads or ∼ 0.5 Terabases. For downstream analysis, we considered all population-wide and genome libraries with mean genome coverage ≥ 100X and ≥ 20X average genome coverage, respectively.

### Genome assembly of isolates

Individual read library alignment files were merged with *samtools v1.8* and were quality checked using *fastQC v0.12.1*. Adapters were trimmed and phiX contamination was removed using *BBDuk* (in BBMap v39.01) to a minimum of 50 bp in length. Reads were k-trimmed from the right with a kmer size of 21, minimum kmer size of 11 and hamming distance of two along with the “tpe” and “tbo” options. Quality trimming was performed from the right with a Q-score of 15. *BBMap*’s reformat.sh was used to interleave read libraries and libraries were merged. The interleaved read library was assembled using *SPAdes v3.15.1* in “isolate” mode with kmers set to 21, 31, 41, 51, 61, 71, 81, 91, 101, 111, 121. BBMap’s reformat.sh was used to remove contigs under 1000 bp from the assembly. The assemblies were binned into putative genomes using *MetaBAT2 v2.15*. Coverage information was calculated using *BBMap* with a 98% identity and the resulting alignment file was sorted using *samtools v1.16.1*. *Metabat2* was run with a minimum length of 1500 bp and default workflow was used in *CONCOCT v1.1.0*. Only the sample’s reads were used to provide coverage information. Putative genomes were also binned using *Metabat2* with no coverage information. A quality comparison of the original assembly and all putative MAGs was performed using *checkM v1.2.2* lineage workflow, alongside coverage information from *BBMap* (at 95% identity). If the assembly was determined to be low contamination (&10%), it was used in later analysis. If the assembly was too contaminated, a MAG of high quality and high coverage (representing the dominant isolate in the sample) was chosen for later analysis. The assemblies and MAGs chosen were annotated using *prokka v1.14.6*.

### Polymorphism detection

For polymorphism detection, we built a custom *python* pipeline, which was developed around *breseq v38.1* (69), a tool designed for polymorphism identification from evolution experiments. We removed adapter sequences and quality filtered raw sequencing files with *fastp v0.23.2* (70), using the parameters below: *-q 20--detect_adapter_for_pe-l 80--cut_tail--cut_tail_window_size 1.* Only polymorphisms supported by at least two forward and two reverse reads were retained downstream in the analysis. In *breseq*, calls were made in polymorphism mode for population samples, while in consensus mode for the isolates. All calls were made relative to the RefSeq genome for the *B. thetaiotaomicron* VPI-5482 strain (accession number; *NC_004663.1*) and plasmid (accession number; *NC_004703.1*). For population-wide samples, we filtered out any genomic sites supported by less than 100 reads and detected with frequency lower than

0.01 in the population. We additionally removed polymorphisms nearly fixed (≥ 0.9 frequency) in the ancestral population that was used for the inoculation of specimens in each experiment. Last, we masked all polymorphisms in regions with detected phase variation, due to the uncertainty that they would introduce false positive calls. For genome annotation, we used all available data from NCBI’s RefSeq database (71) that has been produced with the NCBI Prokaryotic Genome Annotation Pipeline. Additionally, data for COGs were incorporated from the COG database (update 2020) (72). All genome and annotation files can be accessed here: <u>RefSeq Link</u>

### Phase variation detection and masking

Typically, phase variation is characterized by inversions or recombination events between highly homologous genomic fragments that often include inverted repeats, a phenomenon which can mislead SNP calling algorithms into generating false positive calls. However, these polymorphisms would appear or disappear together, depending on the phase. Therefore, in order to identify such genomic signatures, we searched for polymorphisms with highly correlated frequency trajectories, within a relatively small genomic window (100 bp). To verify phase-variable variants, we assembled 170 genomes from isolated colonies (described above). 154 assemblies passed our quality criterion of *N50* > 100K. Assemblies were initially annotated with *prodigal* v2.6.3 (73) and genes were associated to the reference genome according to their best *blastn* hits using *BLAST v.2.12.0+* (74). Then, each contig’s orientation was determined relative to the reference genome, by evaluating the concordance fraction between all genes on the contig and the respective genes on the reference genome. In cases where concordance was < 50%, contig orientation was reversed. We defined phase variation regions (or shufflons), as loci where at least one recombination event would be supported from the genomic data, after manual inspection of all regions associated with a signature of phase variation, considering only cases where all genes involved were physically linked (present on the same contig). *Pfam* protein domains for the N-terminal regions of SusC-family genes were identified through the InterPro database (75) and predicted signal peptides according to SignalP 6.0 (76).

### Enrichment analysis for functional groups

In order to assess whether genetic diversity was distributed unevenly across functional groups, we estimated an enrichment score, accounting both for the size of each group as well as the total number of polymorphisms in each sample. We then applied a *one-sample Wilcoxon test* to evaluate statistical significance of our results from a null distribution with a mean enrichment score of 1. For each *group j* and each sample *i* the enrichment score was given by the formula below:

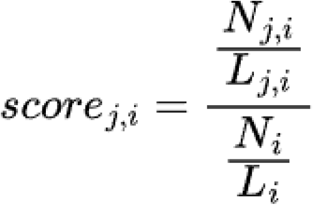

where *N* is the number of polymorphisms and *L* is the length in base pairs. We estimated enrichment scores at the COG category, COG ID and gene level. It is worth mentioning that by definition some genes might be associated with more than one COG family or COG ID. In cases of redundancy, each polymorphism was counted once for each category. Specifically, for the gene level analysis, we only considered cases of statistical enrichment where at least five polymorphisms would be present in a sample.

### Early and late response polymorphisms

Polymorphisms were grouped into early response (ER), late response (LR) and sporadic (SP) based on their prevalence and persistence across evolutionary trajectories. ER polymorphisms should be detected in 4 out 5 samples in at least three separate mouse trajectories. LR polymorphisms should appear either in the last or in the last two sampled time points in at least three separate mouse trajectories. The remaining polymorphisms were considered sporadic.

### Polymorphism association to intestinal compartments

In order to identify strong associations between polymorphisms and intestinal compartments, we used *Maaslin2* (77) to fit a Multivariate Generalized Linear Model (GLM). As input, we used read counts supporting each polymorphism after normalizing with the sequencing library size. We removed any polymorphism with prevalence < 0.2 and also very highly correlated (Pearson r > 0.9) to reduce overfitting. We additionally applied a Mann-Whitney statistical test and adjusted p-values with the false discovery rate (FDR) method. Only consensus significant calls from both approaches were considered strong associations.

### Other statistical tests

We used *compare_means()* function from *ggpubr v.0.6.0* package (78) in *R,* to compare means between or across groups of samples (Wilcoxon signed-rank and Kruskal-Wallis tests) as well as *scipy.stats.wilcoxon* function (79) in *python*, respectively. In paired tests, we did not consider pairs in the calculation where a sample would be missing. For PERMANOVA, we used the *adonis2* function from the *vegan v.2.6-6.1* package (80) in *R*.

### Phase haplotypes identification in complete RefSeq genomes

To evaluate the presence of observed structural variants in other *B. thetaiotaomicron* strains, we used 19 complete assemblies from the RefSeq database available in March 2024 (Table S1). Pangenome was generated using *mmseqs2* (81) under *Panacota v.1.4.2* environment (82) with 60% protein similarity threshold. The phylogenetic tree was obtained based on the concatenation of alignments of single-copy common genes using *IQTree v.2.2.0.3* (83) with 1000 bootstraps and the GTR DNA substitution model. Locally collinear blocks (LCBs) were constructed with *SibeliaZ v.1.2.5* (84), the minimal length of blocks was set to 500bp to reveal structural variants in phase-variable regions. Optimization of parameters for LCB reconstruction and their functional annotation were performed using the *badlon v.0.1.3* (Zabelin et. al. Preprint). Orientation and order of LCBs in the genomes were extracted using *PaReBrick* v.0.5.5 (85) with manual curation of predictions using pan-genome orthogroups.

## Data availability

All sequencing data presented in this study are available online at NCBI’s Sequence Read Archive (SRA) here: <u>PRJNA1277556</u>. Code developed for this study is accessible here: <u>github_repository.</u>

## Acknowledgements

This work was funded by the Austrian Science Fund (FWF; Projects 10.55776/P27831, 10.55776/COE7). The computational results of this work have been achieved using the Life Science Compute Cluster (LiSC) of the University of Vienna. Sequencing was performed by the Next Generation Sequencing Facility at the Vienna Biocenter Core Facilities (VBCF) and the Biomedical Sequencing Facility (BSF) at the Research Center for Molecular Medicine of the Austrian Academy of Sciences (CeMM) in Vienna, Austria.

This study was supported by the Deutsche Forschungsgemeinschaft (DFG) Research Unit FOR5042 (subproject P7) and Collaborative Research Center 1182 (subproject A2) to JFB. The work of O. Bochkareva was supported by the Austrian Science Fund (ESP 253-B), the work of E. Kolodyazhnaya was supported by the RSF grant 24-14-00276.

## Supplementary Information

**Fig. S1.**
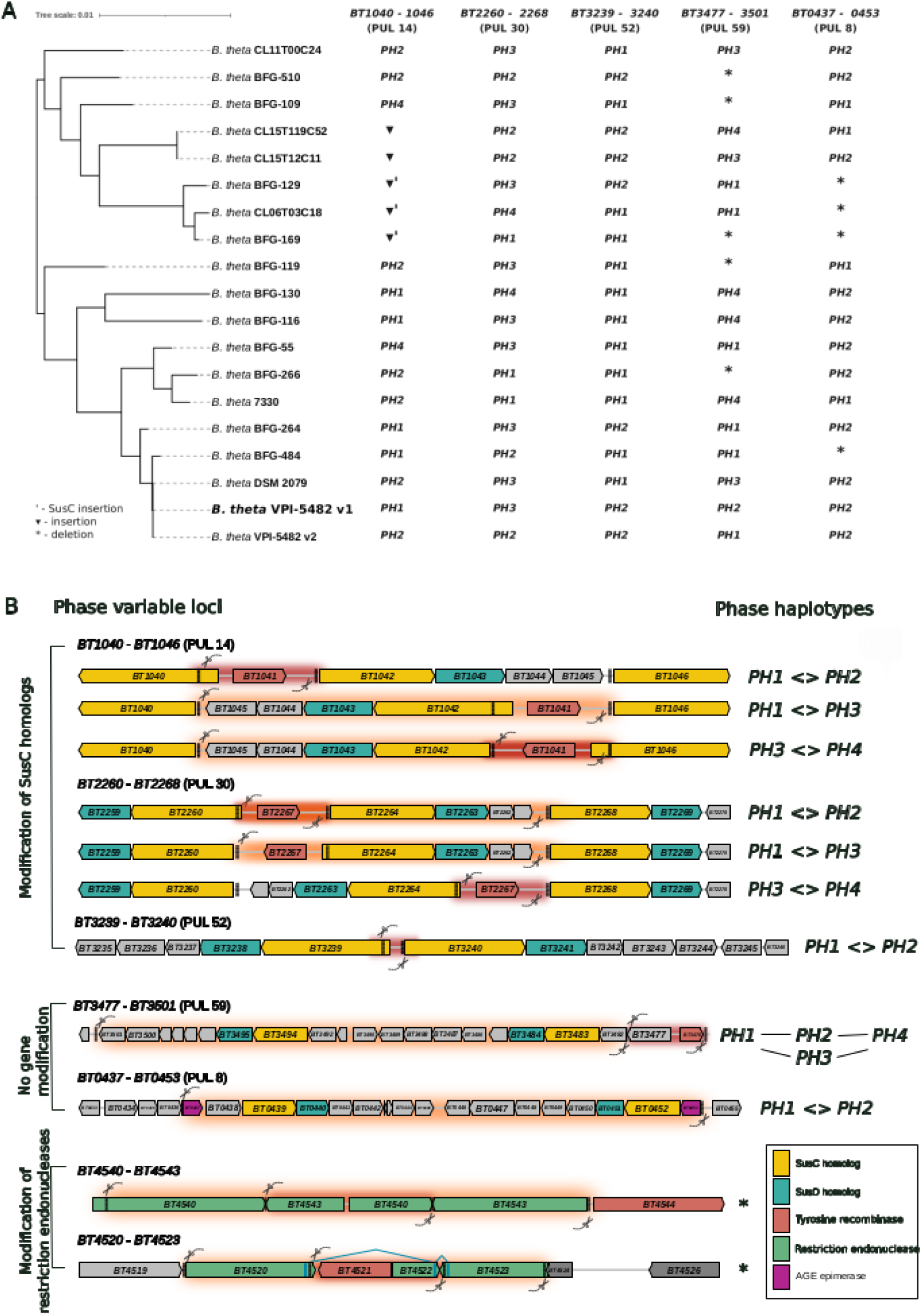
A) Phase haplotypes in 19 Refseq *B. thetaiotaomicron* strains. The distribution of the phase haplotypes observed in complete genomes of *B. thetaiotaomicron* available in RefSeq. **B)** Representation of the types of phase variation detected for each locus.

